# Sequence-to-graph alignment based copy number calling using a network flow formulation

**DOI:** 10.1101/2025.11.21.689771

**Authors:** Hugo Magalhães, Jonas Weber, Gunnar W. Klau, Tobias Marschall, Timofey Prodanov

## Abstract

Variation of sequence copy number (CN) between individuals can be associated with phenotypical differences. Consequently, CN calling is an important step for disease association and identification, as well as for genome assembly validation. Traditionally, CN calling is done by mapping sequencing reads to a linear reference genome and estimating the CN from the observed read depth. This approach, however, is significantly hampered by sequences and rearrangements not present in a linear reference genome; at the same time simple CN prediction for individual graph nodes does not make use of the graph topology and can lead to inconsistent results. To address these issues, we propose Floco, a method for CN calling with respect to a genome graph using a network flow formulation. Given a graph and alignments against that graph, we calculate raw CN probabilities for every graph node based on the Negative Binomial distribution and the base pair coverage across the node, and then use integer linear programming to compute the CN flow through the whole graph. We tested this approach on 15 aligned datasets, involving three different graphs, as well as HiFi and ONT sequencing reads and linear assemblies split into reads. These results demonstrate that the addition of the network flow formulation increases the accuracy of CN predictions by up to 43% when compared with read depth based estimation alone. Additionally, we observed that concordance between predictions from the three different sequence sources was able to reach 93.2%. Floco fills a gap in CN calling tools specifically designed for genome graphs.

## Introduction

Genome sequences often contain copy number (CN) variable segments. This variation can produce impacting differences in terms of molecular behavior, which can ultimately be associated with pheno-typical differences [1–4]. For example, CN changes at the *STRC* locus are known to cause hearing loss in humans [2] and a duplication of two exons in the *Dock2* gene have been associated with striking immune phenotypes in mice [4]. Variation in CN can be observed either within the same sequence or haplotype, or alternatively between sequences or haplotypes. Consequently, CN calling is an important step for disease association and identification, as well as in population genetics. It also appears as a task in genome assembly workflows to determine how often each node in an assembly graph is present in the underlying genome.

Traditionally, CN calling is done by mapping sequencing reads to a linear reference and then estimating the CN in the sequenced sample either from the observed read depth [5–7], by using information from paired-ends and split-read alignments [8, 9], or by combining both approaches [10]. Although these approaches work well within their context, there are inherent problems with using a single linear reference, the biggest of them being the bias against sequences and genomic rearrangements not represented in the linear reference. Moreover, any single reference genome is bound to include numerous unreliable paralogous sequence variants (PSVs)—polymorphic positions where repeat copies differ in the reference genome, but not necessarily in the examined individual, leading to biased read mapping and subsequent CN estimation [11, 12].

In recent years, hundreds of high quality diploid human whole genome assemblies became available, giving rise to several pangenome references [13–16]. Pangenomes reduce the reference bias by using a larger and more diverse set of sequences as reference, increasing the representativity. CN calling can then be done on a bigger set of sequences by using graphs, one of the most common data structures for representing pangenomes, together with sequence-to-graph alignments of sequencing reads. Graphs contain more sequences, some of them completely missing in the linear reference, which then become available for CN estimation. Additionally, graphs also contain non-reference edges between sequences that are not observed next to each other in a linear reference, allowing for a more sensitive read placement, and correspondingly more sensitive CN analysis. The usage of graphs does not change the fundamental reasoning, which is the same as for coverage-based approaches on linear references, where the observed read depth is used to estimate CN, but in this case for each of the graph nodes.

However, using a graph also raises new challenges, in this particular case, the CN consistency across the graph is not guaranteed. That is, CN values are only consistent across the graph if the individual nodes’ CN can be translated into walks through the graph. Several factors can cause erroneous coverage values, including sequencing errors, misalignments and misassemblies, which in turn will negatively impact the coverage-based CN assignments and potentially make them inconsistent across the graph.

To address this problem, we propose Floco, a method for CN calling with respect to a graph (pan)genome using a network flow formulation. Medvedev and Brudno [17] proposed a method to model DNA double strandedness in the context of genome assembly. They developed a framework that, given a set of sequencing reads, builds a bidirected overlap graph from those reads, and estimates read copy counts using a convex min-cost flow formulation. Here, we also use a similar flow-based formulation that, in our case, models node coverages resulting from sequence-to-graph alignments with a negative binomial distribution. In this way, CN assignments are made more consistent throughout the graph compared to estimating CN individually for each node, overcoming incorrect assignments originating from coverage-related noise.

## Method

### Overview

Floco calls individual node CN on genome graphs. As input, it takes a graph (in GFA format) and alignments (in GAF format) against the same graph (Fig. 1a). It then computes the individual base pair coverage for every node and uses them to calculate node CN probabilities (Fig. 1b). The CN probabilities are then provided to our network flow formulation (Fig. 1c), together with some other constraints (Fig. 1d), which we then solve using Integer Linear Programming (ILP). With this method, Floco is able to correct wrong CN assignments made exclusively from coverage-based information (Fig. 1e).

**Figure 1.**
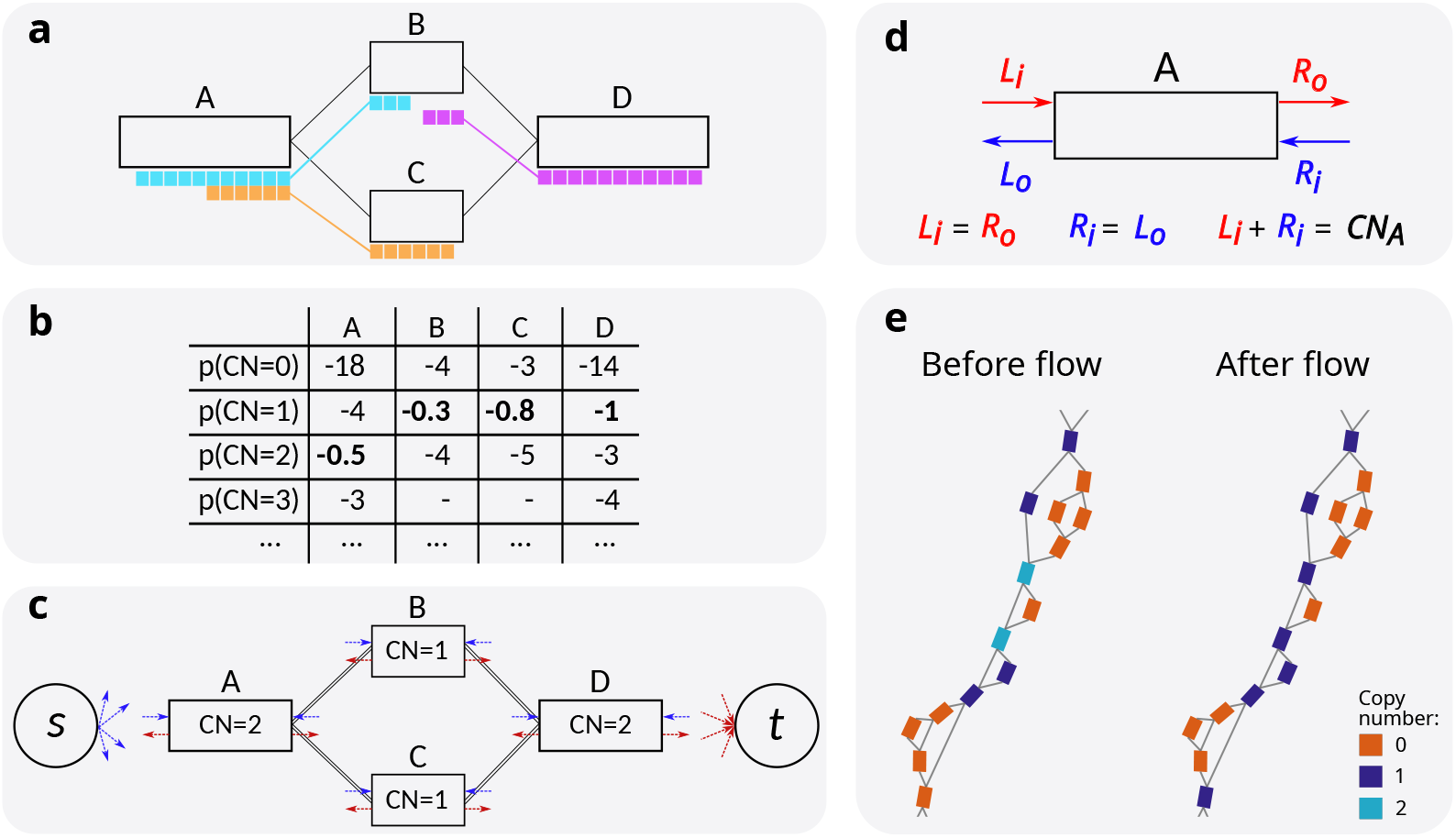
Method overview. **a**, Genome graph and reads aligned against the graph. Reads are aligned to multiple nodes and are shown with different colors. **b**, Table with log CN probabilities per node based on the observed read depth. **c**, Flow network definition and estimated CN. Each connected pair of nodes are linked by two opposite-directed edges. In addition, each node end has one incoming edge from the supersource (denoted by *s*, edge in blue) and one outgoing edge to the supersink (denoted by *t*, edge in red). **d**, Individual node constraints for the flow network: incoming flow on one node end equals to the outgoing flow on the opposite node end; and the sum of incoming/outgoing flow equals to the CN of the node. **e**, strangepg [18] visualization of the CN predictions on a subgraph before and after applying the flow network. On the left side, CN predictions are inconsistent, with an interruption in the path at the bottom of the panel, where all nodes have CN=0 (orange); as well as two diploid nodes (light blue), located in the middle of the image. On the right side, a consistent walk through the graph can be achieved by using nodes with CN=1 (purple).

### Network flow solved with Integer Linear Programming

In order to get our final CN estimations for each node, we use a network flow formulation solved using Integer Linear Programming (ILP) [19]. Minimum cost flow [20] through a graph is a well known problem in computer science. Such flow can be found using a system of linear equations, where *f*_*u𝓋*_ represents flow between nodes *u* and *𝓋*, both belonging to a set of nodes *V* and connected by an edge *e* from a set of edges *E*. The incoming flow through one side of the node *𝓋* equals to the outgoing flow through the opposite side: 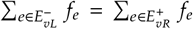, where 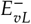 and 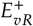 are the sets of edges entering the node *𝓋* from the left and leaving it from the right, respectively (Fig. 1d). Flow through each edge *e* is linearly penalized through costs ξ_*e*_. With additional constraints on the node/edge capacity, as well as on the total required flow, the problem is then defined as the minimization of the total cost of the flow from a source *s* to a sink *t*: ∑_*e*∈*E*_ *f*_*e*_ · ξ_*e*_ → min.

With regards to the CN flow, we adapt the minimum cost flow formulation into a maximum probability flow on a bidirected graph, where each edge *e* has integer flow *f*_*e*_ and each node *𝓋* has associated probabilities *p*_*𝓋c*_ (Equation 2) as well as a range of possible CNs *C*_*𝓋*_. CN is encoded with a non-negative integer variable *x*_*𝓋*_ that is connected to CN probabilities via a piecewise linear (PWL) function *g*(*x*_*𝓋*_; *p*_*𝓋*_), where *g*(*x*_*𝓋*_ = *c*; *p*_*𝓋*_) is set to log *p*_*𝓋c*_. We also extend the concept of source and sink to a supersource *s* and a supersink *t*, each of them connected to both ends of every node *𝓋* ∈ *V* through a directed edge, with *s* and *t* having only outgoing and incoming edges, respectively. The usage of such edges is subject to a “cheap” cost if the node has no other edges connected to the same node end (−25 by default); or “expensive” cost otherwise (−10^5^ by default); corresponding values are stored in the edge costs ξ_*e*_. The impact of different edge costs on the CN calling accuracy can be found in the Suppl. Fig. 3.

In addition, we apply a cost on the usage of edges supported by few reads. Specifically, we examine the number of reads *d*_*e*_ supporting the edge *e*, i.e., the number of reads consecutively passing both of the edge nodes; and compare *d* with the expected number of reads *E*[*d*_*e*_] given the edge length *l*_*e*_ (size of the overlap between the two nodes). We have observed that read length values follow a skew normal distribution (SN) [21] (Suppl. Fig. 1), and calculated *E*[*d*_*e*_] as *S*_SN_(*l*_*e*_) · *μ*_1_, where *S*(*l*) is the SN survival function of the edge length (expected number of reads longer than *l*_*e*_) and *μ*_1_ is the expected read depth for nodes of length 1, defined later. Edge *e* remains free whenever *d*_*e*_ ≥ *E*[*d*_*e*_]/4, otherwise we set edge cost ξ_*e*_ to the “cheap” superedge penalty. In summary, CN flow is formulated as the following maximization problem:

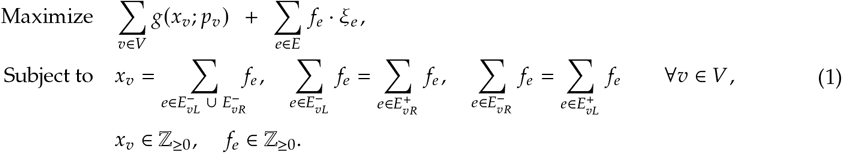

Finally, Floco employs Gurobi ILP solver [22] to find the most likely set of CNs *x*.

### Node coverage computation and background read depth

As nodes in the overlap graphs contain redundant sequence, which could influence observation independence, Floco first clips nodes in a procedure similar, but less strict than graph bluntification [23] (see Suppl. Methods). We then define node coverage as the sum read depth across all retained positions in the node, obtained by summing up alignment sizes between reads and clipped nodes for all primary alignments in the input GAF file. Fully clipped nodes are not considered for the coverage computation, and their CN is estimated purely based on the flow. In addition, we split all nodes longer than *m* (100 bp by default) into bins of that size. Bin coverages are defined and calculated analogous to the node coverages.

In previous studies, negative binomial (NB) has been used to model the total number of reads overlapping a single genomic position or a fixed-size bin [11, 24]. In contrast, we defined coverage as the sum read depth over all base pairs in a given node or bin as it provides a better ability to capture reads, stopping or starting within a node/bin, as well as allows for variable length nodes shorter than the bin size *m*. Nevertheless, we empirically observed that node/bin coverage can also be explained by the NB distribution (Suppl. Fig. 2a-b). This way, when bin coverage for nodes with CN=1 follows NB(*r, q*), bin coverage at CN=*c* follows NB(*cr, q*) due to the additive characteristics of the NB distribution. At CN=0 a node should not be covered by any reads, nevertheless, non-zero coverage can be observed due to read alignment errors. We model that using discretized exponential distribution as it has a lower tail than NB distribution with small *r*. To connect NB and exponential parameters, we set exponential mean to *ε* · mean(NB), resulting in 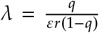, where *ε* represents putative read aligner error rate (default: 0.02). Finally, we performed χ^2^ goodness of fit test and confirmed that the mixure of exponential and NB distributions explains observed bin coverages well (P value ≈ 1).

To estimate NB parameters, we examine bin coverage across nodes with at least 10 bins as longer nodes are expected to have better read alignments. Next, we discard outliers as well as potentially uncovered nodes by first removing nodes where all bin coverages are under *m* (effectively, no single read passes through the whole bin) and then removing the top and bottom 3% of the nodes based on their mean bin coverage. This is done to reduce the number of variables and simplify numerical optimization due to extremely high CN of some nodes, as well large number of completely uncovered nodes. After node filtering we estimate coverage parameters using the maximum likelihood method by numerically optimizing the total log-likelihood of the retained bin coverages under a weighted mixture of distributions for CNs from 0 to *c*_max_ = *c*_def_ + 2, where *c*_def_ denotes the expected CN for most nodes in the graph. By default, Floco tests *c*_def_ = 1 and 2 and selects one that produces higher log-likelihood. In addition, we allow users to manually set *c*_def_ due to the fact that in certain cases a graph consisting of mostly CN=1 nodes is almost indistinguishable from a graph containing mostly CN=2 nodes. Optimization is performed using the bounded Nelder-Mead method [25, 26] on the following variables: CN=1 distribution mean and variance; and fractions of node CNs, restricted between 0 and 0.45 for *c* ≠ *c*_def_ and between 0.5 and 1 for *c*_def_. For better conversion, node CN fractions are normalized before the likelihood calculation instead of numerical optimization using one less variable.

### Node coverage probabilities

Computed background read depth distribution with mean 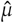 and standard deviation 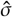 describes a coverage for a bin of size *m* and CN=1. To extend this definition to variable size nodes, we empirically observed that the distribution of mean and standard deviation proportionally depend on the bin size: 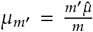 and 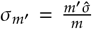 (Suppl. Fig. 2b). This allows us to define read depth distribution NB(*r*_*𝓋*_, *q*_*𝓋*_) for any node *𝓋* with CN=1 using NB method of moments. As above, we define probability of observing read coverage *k*_*𝓋*_ at CN *c* > 0 as *P*(*k*_*𝓋*_ | *c*_*𝓋*_=*c*) = *P*_NB_(*k*_*𝓋*_; *cr*_*𝓋*_, *q*_*𝓋*_) and use the exponential distribution for CN=0. To improve predictive power of large nodes and preserve independence of observations, we randomly subsample node bins so that on average each read appears in exactly one bin (rate = *m* / mean read length). Whenever less than two bins remain per node, we use node coverage for calculating CN probabilities, otherwise we redefine coverage probability as the product of all bin coverage probabilities: 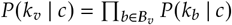, where *B*_*𝓋*_ is the set of retained node bins.

Afterwards, we use Bayes’ theorem with equal priors to define bin or node CN probabilities based on the observed coverage *k*_*𝓋*_:

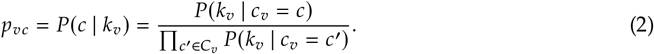

To define the set of possible CNs *C*_*𝓋*_ for node *𝓋* we start from normalized and rounded coverage 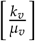 and extend the set to the left and right until log-difference between best and current probability log *P*(*k*_*𝓋*_ | *c*_best_) − log *P*(*k*_*𝓋*_ | *c*) does not get too large (over 4 expensive superedges).

## Results

To evaluate Floco and demonstrate its utility, we focus on two applications: The first application is *assembly graphs quality control (QC)*, where we align reads from a sample against its own assembly graph. In this case, we expect all nodes to have CN>0, given they were derived from reads from that same sample, while nodes with CN=0 indicate errors. CN calling on assembly graphs also has applications for phasing workflows [27, 28], but here we restrict our attention to QC.

The second application is *CN calling using a pangenome reference*, where we align reads to a pangenome graph. When aligning reads from a sample that was used during graph construction (in-sample), the expected outcome is that all nodes originated from that sequence are covered and all others are not. When aligning a sample that was not included in graph construction (out-of-sample), it potentially carries yet unobserved CN variation, which can be reflected in an increase in the number of nodes with CN=0, indicating deleted sequence, or nodes with CN>2, indicating duplicated sequence.

In our evaluations, we contrast the CN assignment *before* and *after* using the network flow. That is, *before flow* corresponds to using the same negative binomial formulation to assign a maximum likelihood CN to each node separately without considering the graph connectivity, where *after flow* corresponds to the full Floco approach.

### Data description

We used three graphs for evaluation (Suppl. Table 1): For the assembly QC use case, we used two Verkko [28] assembly graphs, one from a diploid human HG01114 [16] (4k nodes and 3.4k edges) and the other one from a tetraploid potato [29] (20k nodes and 27k edges). For the pangenome use case, we used the minigraph [30] pangenome graph from the Human Pangenome Reference Consortium (HPRC) [15], which contains 90 haplotypes, being the most fragmented of the graphs (671k nodes and 965k edges).

Furthermore, we used the following sequencing and synthetic data (Suppl. Table 2), aligned to the graphs using GraphAligner [31] (Suppl. Methods). First, we aligned HG01114 and Altus sequencing data to the corresponding assembly graphs. Next, we aligned three samples to the HPRC pangenome graph: haploid CHM13 [32], which was used as the graph’s backbone sequence; HG01258, which was added to the graph during its construction (in-sample); and HG01114, which is not present in the graph (out-of-sample). We use suffixes ‘asm’ (assembly graph) and ‘pang’ (HPRC pangenome) to indicate which graph was used. This way, we consider five sample–graph pairs: HG01114-asm, Altus-asm, CHM13-pang, HG01258-pang, and HG01114-pang (Table 2). For each sample we used PacBio High Fidelity (HiFi) and Oxford Nanopore (ONT) reads, as well as their own linear assemblies, “chopped” into groups of fixed-size reads (see Suppl. Methods).

**Table 1.**
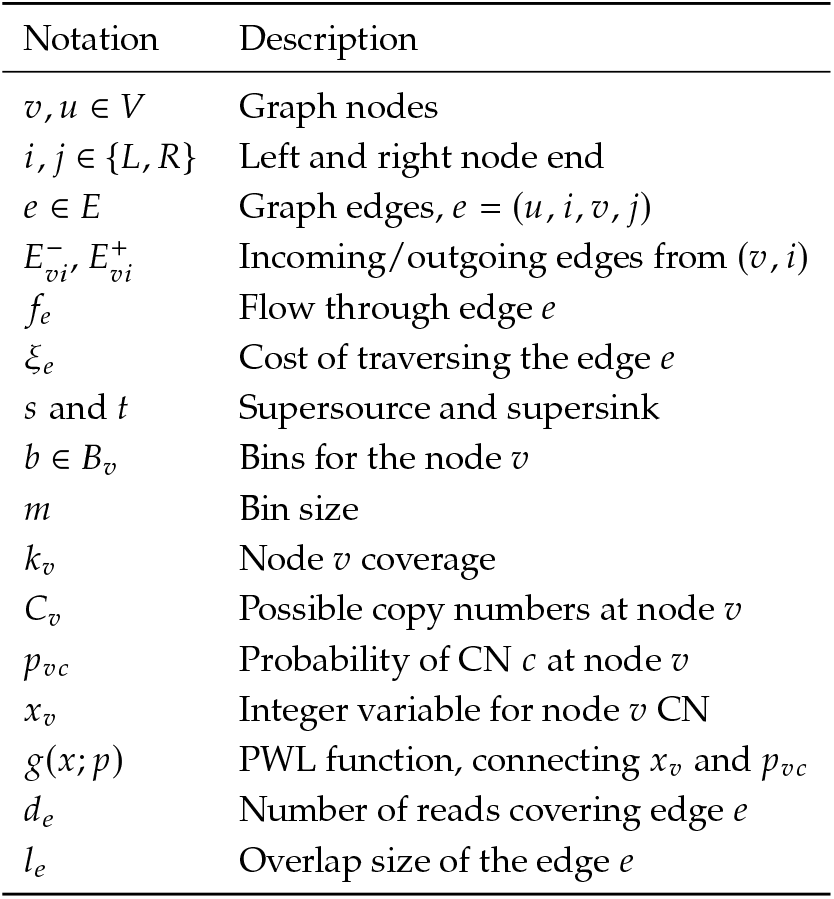
Table of notation.

**Table 2.** Sample–graph pairs used for Floco evaluation. First column shows the sample–graph pair, where the graph is abbreviated as either ‘pang’ (HPRC pangenome) or ‘asm’ (assembly graph). The last column (“in-sample”) defines whether the sample was used for the graph construction.

| Pair | Species | Graph | In-sample |
| --- | --- | --- | --- |
| CHM13-pang | *Homo sapiens* | HPRC | Yes |
| HG01258-pang | *Homo sapiens* | HPRC | Yes |
| HG01114-pang | *Homo sapiens* | HPRC | No |
| HG01114-asm | *Homo sapiens* | HG01114 | Yes |
| Altus-asm | *Solanum tuberosum* | Altus | Yes |

### Floco accurately calls copy number for CHM13

CHM13 was used as a backbone when constructing the HPRC graph [15] and mapping CHM13 reads to the HPRC graph (CHM13-pang) hence constitutes an in-sample setting. For the ground truth CN set, we defined all nodes belonging to CHM13 (identified by the SN tag in the rGFA file) as having CN=1 (314k nodes), whereas all other nodes were assigned CN=0 (357k nodes). We then compared the CN predictions from the CHM13-pang chopped assembly, HiFi and ONT alignments against the ground truth, both before and after applying the network flow. We then measured accuracy as the fraction of nodes with correctly estimated CN as well as the fraction of total sequence, represented by these nodes.Additionally, we evaluated the effect of read depth on CN prediction accuracy by downsampling sequencing data to coverages from 1× to 30×.

First, we observed that Floco achieved almost perfect accuracy on the chopped assembly (Fig. 2, Suppl. Fig. 4), accurately predicting CNs for 99.7% (99.6% before flow) of the nodes, and achieving perfect precision (error <4·10^−5^). When translated into the sum of node lengths, around 1.9% of the total sequence was predicted to have CN=0 instead of CN=1, driven primarily by the missing read alignments. Even then, using the network flow allowed to correctly predict CNs for additional 619 Mb compared to CN predictions before the flow. So while a limited amount of residual errors remain, this demonstrates that Floco effectively leverage the graph topology to derive CN assignments.

**Figure 2.**
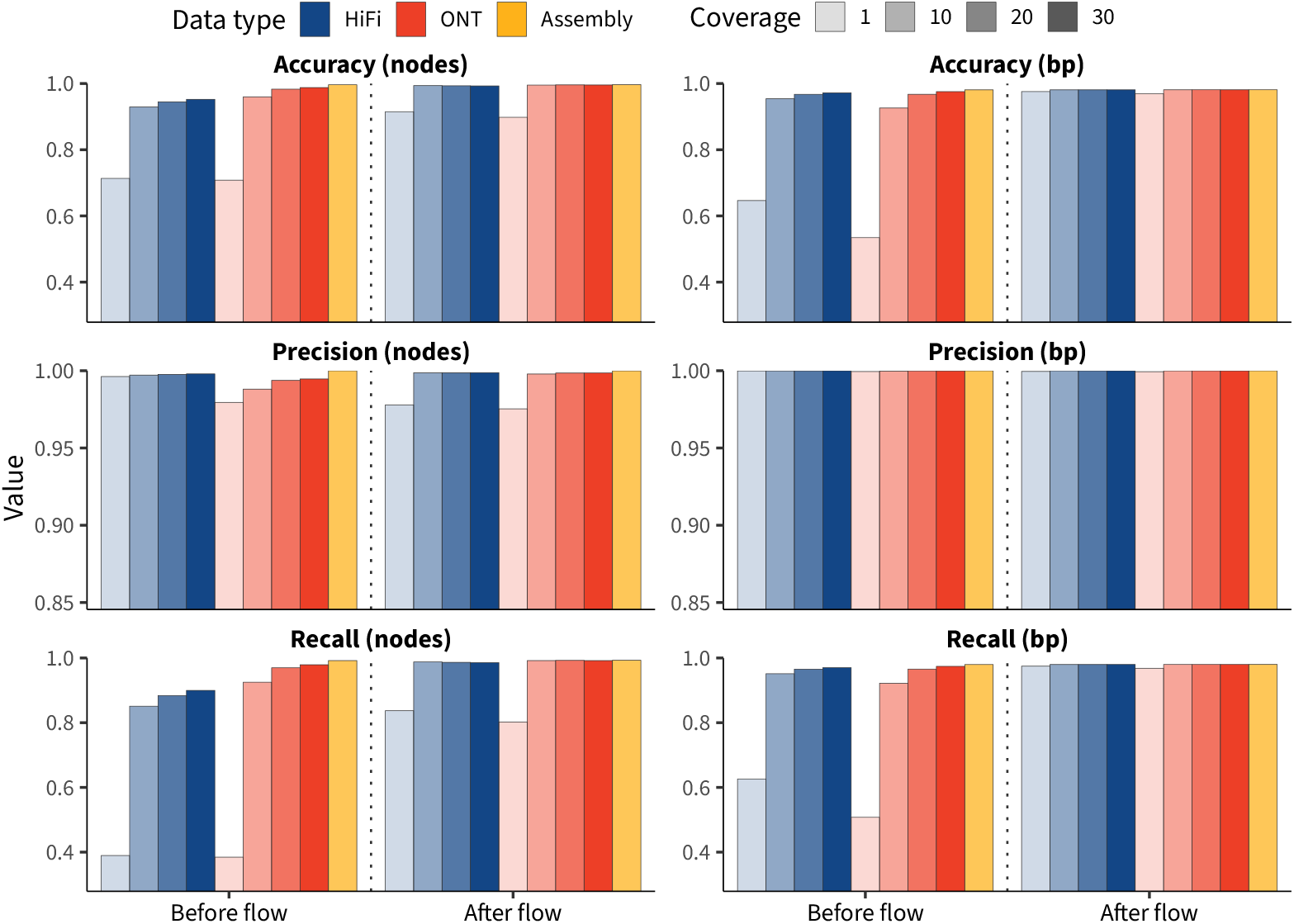
Copy number calling accuracy for the CHM13 sequencing data and HPRC pangenome graph. Each subpanel shows prediction evaluation before and after CN flow, calculated for the PacBio HiFi (blue), Oxford Nanopore (red) and the chopped assembly (yellow) datasets, subsampled to coverages between 1 and 30 (shown with different transparency levels). Left and right groups of panels show values calculated based on the number of nodes and the sequence length, respectively. Accuracy is either measured directly (fraction of nodes/sequence length matching with the CHM13 path in the graph); or through corresponding precision and recall. Y-axis is clipped at the bottom at 0.35 for accuracy and recall and at 0.85 for precision.

At 30× coverage, Floco accurately predicted CNs for 98.1% of the total sequence for both the ONT and HiFi data, trailing behind assembly-based CN predictions by just 0.007–0.016 percentage points. At the same time, CN predictions without the network flow achieved 97.5% (ONT) and 97.2% (HiFi) accuracy, 0.6 and 0.9 percentage points behind the flow-based accuracy values. In terms of the number of nodes, improvement was even greater: 0.8 percentage points (98.8→99.6%) for ONT and 4.1 percentage points (95.2%→99.3%) for HiFi sequencing data (Fig. 2).

At smaller coverage it becomes more difficult to both accurately estimate background read depth, as well as to distinguish sequence, unsupported by chance, from sequence truly missing from the sample genome. Consequently, network flow, which effectively aggregates signal from nearby nodes, achieves a greater edge over CN predictions for individual nodes. At 1× coverage data, ONT-based CN predictions showed 53.4% and 96.9% accuracy (total length) before and after the network flow, while HiFi-based accuracy marked 64.6 and 97.6%, respectively, which represents approximately a 15-fold reduction in error rate in both cases. In contrast to other cases, some CN=0 nodes were assigned CN=1 (node-based precision = 97.5–97.8%), however, this mostly affected short nodes (median length ≈70 bp for both sequencing technologies), and did not notably influence precision, weighted by the sequence length.

### Concordance between technologies

Obtaining ground truth CN values is a more difficult problem for other samples and graphs, even in the presence of linear assemblies. We therefore focused our analysis on comparing Floco CN predictions for three different alignment sources: independently generated HiFi and ONT sequencing datasets, as well as sample linear assemblies, chopped into reads.

In the human assembly graph case, for HG01114-asm (Fig. 3a, Suppl. Fig. 5a), Floco predicted the same CN values for all three data sources across 3,956 Mb (93.2% of total sequence in the graph), corresponding to 73.5% of the informative nodes, which contained unique sequence following graph preprocessing (Suppl. Methods). Individually, both HiFi and ONT CN predictions matched the chopped assembly across 96.5% and 96.0% of the total sequence, respectively; as well as matched each other by 93.8% (83.8% nodes). However, we observed that discordances are often caused by Floco selecting one or the other of two alternative paths through a bubble, where both arms have almost identical sequences and very similar read depth (Suppl. Fig. 6). Consequently, while difficult to measure, Floco concordance in terms of the actual sequence content is likely to be significantly higher than the raw sequence length concordance. We also note that such bubble structures (Suppl. Fig. 6) exist only in places where the assembly tool was not able to resolve them; that is, they by construction reflect loci with ambiguity, accumulation of systematic sequencing errors, or artifacts introduced by the assembly algorithm.

**Figure 3.**
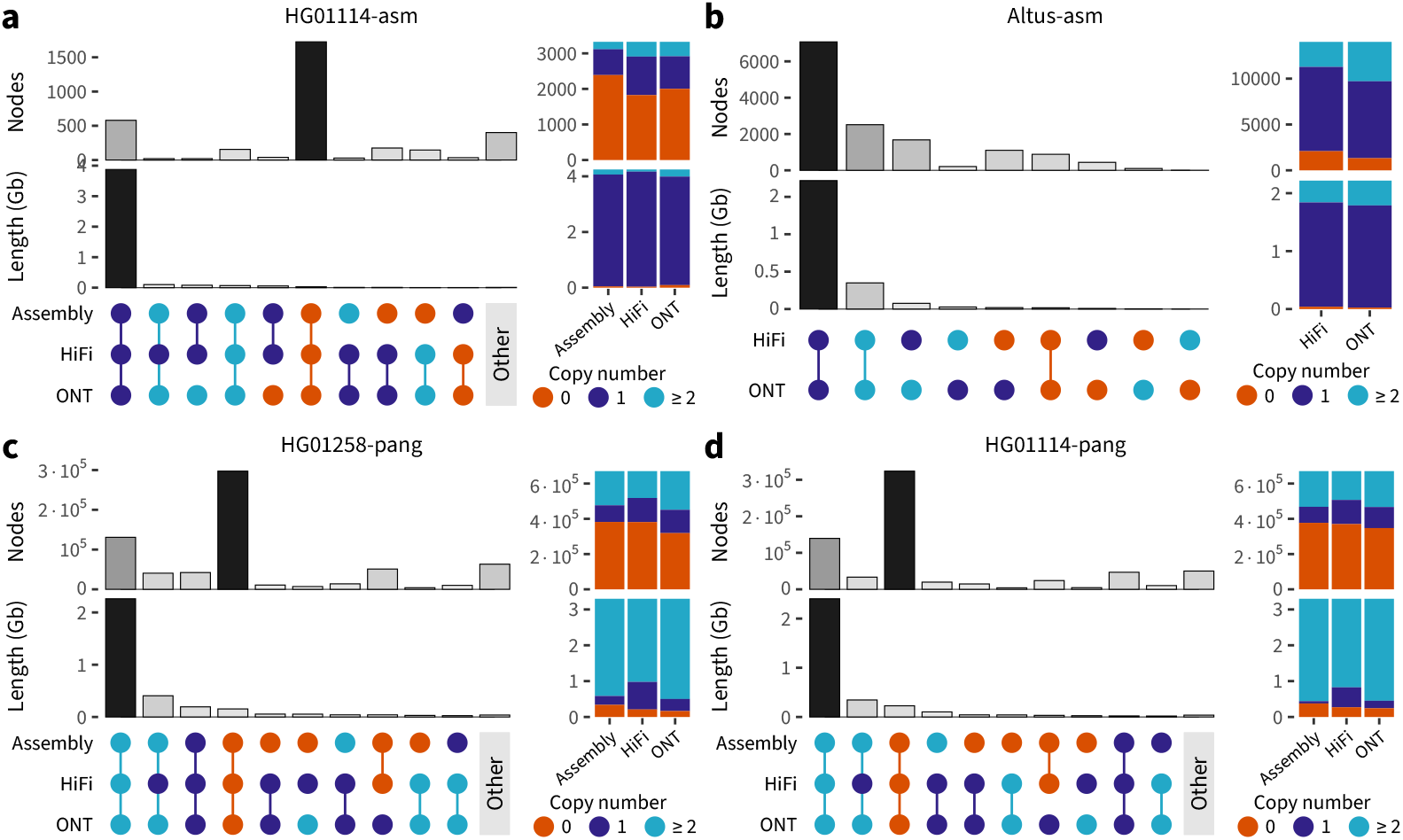
Concordance between copy number predictions for different sequencing technologies. Each of the four panels contain an extended UpSet plot, which shows the number of nodes and sum length of the nodes for each combination of CN predictions (orange for 0; purple for 1; light blue for larger values) based on the chopped assembly, PacBio HiFi and Oxford Nanopore (ONT) reads. Columns are sorted by sum length, 11th and later columns are groupped together (“other”). Right side of each panel shows the total number of nodes and their sum length for each CN group and each sequencing technology. Concordance is calculated for **a**, HG01114 human data and **b**, potato Altus data mapped to their corresponding assembly graphs; as well as **c**, in-sample HG01258 and **d**, out-of-sample HG01114 data mapped to the HPRC pangenome. Chopped assembly is removed from the panel **b** due to the low resolution of the corresponding linear haplotype-resolved assembly.

In the tetraploid potato assembly graph case, Altus-asm (Fig. 3b), we excluded assembly-based predictions, because of low assembly quality (see Suppl. Fig. 5b). For the two sequencing datasets, Floco predicted the same CNs across 92.9% of the sequence (71.4% informative nodes), HiFi and ONT-based CN predictions, reaching concordance levels similar to HG01114-asm.

Next, for the HPRC pangenome graph, we examined in-sample HG01258-pang and out-of-sample HG01114-pang alignments. In both cases, ONT better matched the chopped assembly, with 89.3 and 90.6% of concordant sequence, respectively, compared to HiFi data with 80.7 and 81.4% of concordant sequence. This can be explained by the prevalence of CN=1 sequence at the expense of higher CN in the HiFi-based predictions (Fig. 3c–d, Suppl. Fig. 5c–d). Nevertheless, in terms of the node-based concordance, Floco showed similar values for both technologies: 81.4% (HiFi) and 75.6% (ONT) for HG01258-pang compared to 81.6% and 81.7% for HG01114-pang. Predicted node CNs had high correlation between the two sequencing datasets: *r*=0.833 and 0.899 for HG01258 and HG01114, respectively.

Even though HG01258 was used for the HPRC graph construction, its ONT data has lower coverage (7× compared to 26× for HG01114, see Suppl. Table 3), which could explain lower prediction concordance. Additionally, we looked at the concordance for in-sample CHM13-pang, where the concordance between the three sources reached 99.9% of concordant sequence (99.4% of informative nodes) (Suppl. Fig 4).

#### Assembly graph quality control

With Floco, we are able to flag potentially misassembled nodes with CN=0, even though some of them have observed coverage. For example, when aligning HiFi reads from HG01114 against its assembly graph (HG01114-asm), we get 33.9 Mb of sequence with CN=0, corresponding to 1,825 informative nodes. Most of these nodes belong to very small components of the graph, where alignment is harder to perform. There are, however, some cases in the bigger components (Suppl. Fig. 7), where bubbles are erroneously created, leading to three possible paths through a given locus, when at most two would be the expected for a diploid individual. We have manually examined some of these examples and found that all of them appear to be indeed misassembled nodes, where the sequence content of the node is represented in the alternative paths, either exactly or with very high sequence similarity.

Additionally, we examined CN predictions based on the HG01114 ONT read alignments to the HG01114 assembly graph, and observed the same pattern, although the number of CN=0 nodes was slightly higher in that case (2,208), corresponding to a total of 91.2 Mb of sequence. The difference between the two sequencing technologies is explained by the larger number of CN=0 nodes in smaller components. This is likely due to the fact ONT reads have a higher error rate than HiFi reads, introducing additional noise in the alignment process. We also note that small nodes in these (otherwise high quality) assembly graphs typically represent either errors or low complexity sequence.

#### Runtime and peak memory usage

Floco CN calling on assembly graphs (HG01114-asm and Altus-asm) is extremely lightweight, and can easily be executed on a laptop; with one-threaded runtime varying between 8 and 22 minutes (Suppl. Table 3) and peak memory not exceeding 3.1Gb. As pangenome graphs are more convoluted, Floco required more time (ranging from 40 minutes to just under 3 hours), with peak memory not exceeding 53Gb for real sequencing data. In any run, Floco CN calling took a fraction of time needed to align reads to the graph, as GraphAligner required between 14 and 100 hours, depending on sequencing coverage, for both graph types, and consumed between 8 and 35Gb.

Furthermore, we evaluated the impact of varying coverage on the Floco computational resources usage. Starting from 33× (HiFi) and 112× (ONT) coverage for CHM13-pang, we downsampled the alignments and measured Floco runtime and memory consumption. As Figure 4a shows, running time weakly grew with coverage for both sequencing technologies (Pearson’s *r*=0.83–0.86), with CN calling taking between 79 and 100 minutes for the HiFi data and between 89 and 131 minutes for the ONT data.

**Figure 4.**
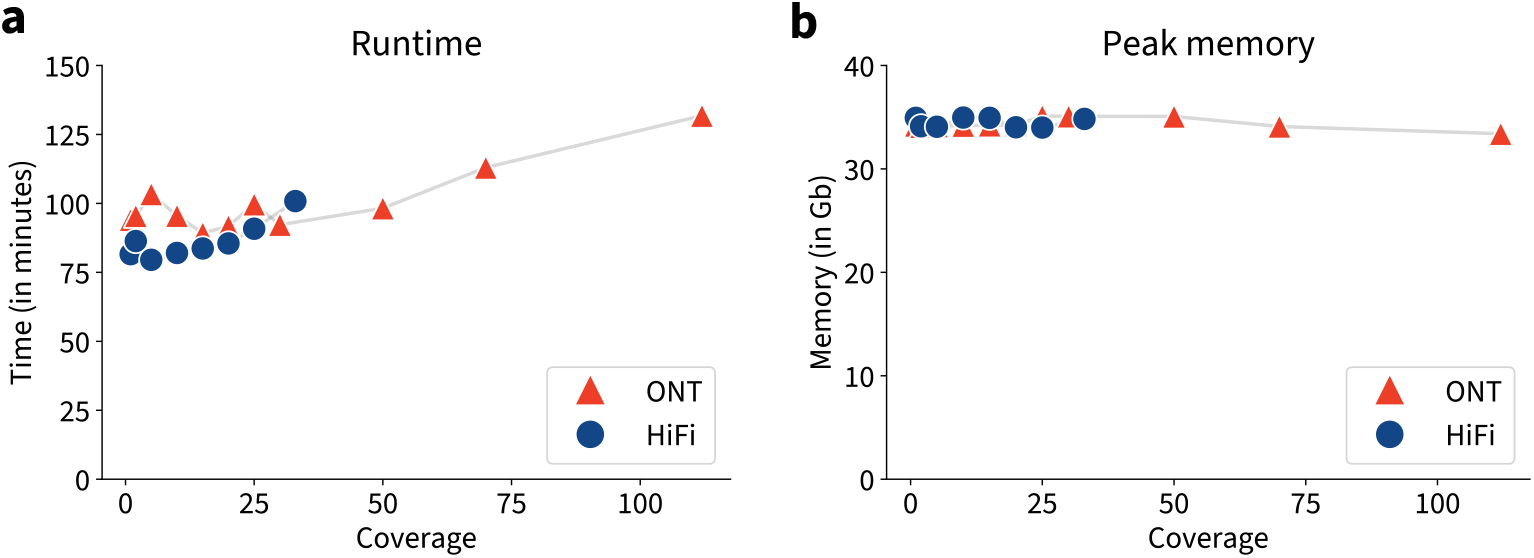
Computational resource usage per sequencing coverage. Floco runtime (**a**) and peak memory usage (**b**) were measured for the ONT (red triangles, coverage between 1 and 112) and HiFi data (blue circles, coverage between 1 and 33).

At the same time, memory usage did not depend on the underlying coverage, with Floco consuming 33.4–35.1Gb in all cases (Fig. 4b).

## Discussion

We presented a new method, Floco, to call CN on genome graphs using graph-aligned reads. Floco combines techniques for read-depth based CN estimation [5–7, 10] and network flow formulations introduced in the context of genome assembly [17], addressing a need in the tool ecosystem for working with genome graphs. Even though the proposed flow formulation involves piecewise linear node weights in addition to linear edge weights, the problem can be reduced to the regular min-cost network flow problem, which can be solved in polynomial time [17, 20]. Floco uses the integer linear programming solver Gurobi [22] to find the optimal flow, which leads to fast runtime on all evaluated datasets. We therefore did not attempt to use specialized flow solvers.

We demonstrate the advantage of using the network flow approach by comparing CN predictions before applying the flow, i.e. using read depth information only, and after applying the flow to correct inconsistencies and overcome misalignments and sequencing errors. Results show a clear gain when adding the network flow formulation on top of the read depth based approach (Fig. 2), especially regarding accuracy and precision.

We also demonstrate that Floco is robust with respect to the choice of input data, as shown by the concordance observed between the predictions made from different sources (Fig. 3). Given the varying characteristics of each source, with ONT reads being much longer than HiFi reads and assembly pieces; while having, on the other hand, a higher error rate; differences in final results are expected. Nevertheless, we still achieved high levels of concordance for at least two out of three sources, reaching values above 92.9% of the total sequence length for both assembly cases, and 80.7—90.6% for the pangenome cases. The difference between species might be explained by the lower quality of both the Altus graph and its linear assembly, as well as the fact that it represents an autotetraploid organism.

Floco is fast and has low memory requirements (Fig. 4, Suppl. Table 3). Computational resource usage can be reduced further by tweaking input parameters, specifically the “expensive” superedge penalty (see Method), which allows to control a trade-off between speed and accuracy. Additionally, ILP solving time can be decreased by using multiple threads (values shown in this study were all ran with a single thread). Time and memory consumption are mostly influenced by the graph complexity, namely the number of nodes and edges, which translate into more variables for the ILP (see Method).

For the analysis performed here, we have used only long reads. Theoretically, the tool should work with any type of reads, but it is dependent on the input alignments of the reads to the graph, which are more challenging to obtain for short reads. While Floco is a generic tool designed to work on as many graphs as possible, the performance in practice and interpretability of results depends on the input graphs. Especially when quantifying CNs relative to a pangenome reference, the quality of results will depend on whether copy-number-variable segments are represented as their own nodes. As future work, we envision to add functionality to call CN changes that occur within single nodes. Note that although at this point Floco has only been tested using variation graphs, it should work with other types of graphs, like de Bruijn graphs, given its abilities to resolve overlaps and start/end the flow anywhere using superedges.

There are not yet community-curated benchmark sets for CN calling on graphs. Here, we have created our own benchmarks by using publicly available resources. One open challenge consists in comparing CN calls on graphs to CN calls with respect to a linear reference genome. Such a comparison would require a transfer of CN predictions from a graph to a linear reference, involving copy-number-variable regions as well as sequence missing from the reference. This operation is not only difficult to define and develop, but would likely introduce its own discordance. We therefore did not include comparisons to CN callers for linear references. We show that further challenges for the evaluation stem from cases of high sequence identity across different genomic paths through the graph, which can lead to underestimated concordance values.

Our method supports multiple use cases. On the one hand, predicting CNs by aligning a linear assembly against the individual’s assembly graph can be used as quality control for the graph itself (Suppl. Fig. 7). On the other hand, Floco fills an important gap for genotyping classes of genomic variation with variable CN, like segmental duplications, which are becoming more tractable given the advances on assembly methods. This use of Floco has applications for downstream association studies to discover correlations between CN states and phenotypes.

In summary, we have presented a method, implemented in the general-purpose tool Floco, that fills a gap in CN calling tools specific to genome graphs, offering an accurate and robust strategy with multiple applications.

## Data Availability

The HPRC graph used for the benchmarking of Floco can be found at s3-us-west-2.amazonaws.com/human-pangenomics/pangenomes/freeze/freeze1/minigraph/hprc-v1.0-minigraph-chm13.gfa.gz. The Altus tetraploid assembly graph can be found at zenodo.org/records/17599056. The HG01114 assembly graph was produced under the scope of HGSVC3 [16]. CHM13 linear assembly can be found in the European Nucleotide Archive with the accession number PRJNA559484, and the reads can be found at github.com/marbl/CHM13/blob/master/Sequencing_data.md (HiFi Data and Guppy 5.0.7). Data for HG01258 can be found at s3-us-west-2.amazonaws.com/human-pangenomics/index.html?prefix=working/HPRC/HG01258. HG01114 reads can be found in the European Nucleotide Archive with accession number SAMEA115605425. The sequencing reads for Altus are available via the NCBI BioProject under accession number PRJNA778192 (HiFi) and PRJNA1049180 (ONT).

## Code Availability

Floco is implemented in the Python programming language. The source code is freely available under the terms of the MIT license at github.com/hugocarmaga/floco along with installation and usage instructions. Benchmarking scripts are available at github.com/hugocarmaga/floco-benchmarking.

## Acknowledgements

This research was supported in part by funding from the National Institutes of Health National Human Genome Research Institute (grant no. U01HG013748 to T.M.) and from funding from the German Research Foundation (grant no. 525152594 to T.M.).

## Notes

### Competing Interest Statement

The authors have declared no competing interest.

### Summary of Updates

Revisions after reviewing process by RECOMB: Added further explanation on some results; Added supplementary figures; Rephrased some unclear statements.

