## Supplementary Material for "Sequence-to-graph alignment based copy number calling using a network flow formulation"

### **Supplementary Information** for Sequence-to-graph alignment based copy number calling using a network flow formulation

Hugo Magalhães [1,2], Jonas Weber [3], Gunnar W. Klau [2,4],
Tobias Marschall [1,2,✉] & Timofey Prodanov [1,2,✉]

[1] Institute for Medical Biometry and Bioinformatics, Medical Faculty, Heinrich Heine University, Düsseldorf, Germany. [2] Center for Digital Medicine, Heinrich Heine University, Düsseldorf, Germany. [3] Institute of Medical Microbiology and Hospital Hygiene, Heinrich Heine University, Düsseldorf, Germany. [4] Algorithmic Bioinformatics, Department of Computer Science, Heinrich Heine University, Düsseldorf, Germany. ✉Corresponding authors: `` and ``.

**1 Supplementary Figures and Tables** **2**

### 1 Supplementary Figures and Tables

**Supplementary Table 1. Detailed information about the graphs used for the analysis.**

| Graph | Species | Graph constructor | Nr. nodes | Nr. edges | Nr. haplotypes | Total length (Mb) |
|---|---|---|---|---|---|---|
| HPRC | *Homo sapiens* | minigraph | 670,576 | 964,816 | 90 | 3,297 |
| HG01114 | *Homo sapiens* | verkko | 4,077 | 3,400 | 2 | 4,284 |
| Altus | *Solanum tuberosum* | verkko | 20,216 | 26,566 | 4 | 2,797 |

**Supplementary Table 2. Detailed information about the samples used for the analysis.**

| Sample | Species | Alignment source | Read N50 |
|---|---|---|---|
| CHM13 | *Homo sapiens* | Assembly | 20,000 |
| CHM13 | *Homo sapiens* | HiFi | 17,781 |
| CHM13 | *Homo sapiens* | ONT | 56,695 |
| HG01258 | *Homo sapiens* | Assembly | 20,000 |
| HG01258 | *Homo sapiens* | HiFi | 20,342 |
| HG01258 | *Homo sapiens* | ONT | 31,858 |
| HG01114 | *Homo sapiens* | Assembly | 20,000 |
| HG01114 | *Homo sapiens* | HiFi | 21,191 |
| HG01114 | *Homo sapiens* | ONT | 140,864 |
| Altus | *Solanum tuberosum* | Assembly | 20,000 |
| Altus | *Solanum tuberosum* | HiFi | 17,748 |
| Altus | *Solanum tuberosum* | ONT | 25,095 |

**Supplementary Table 3. Floco runs.** Left-to-right, columns describes sample-graph pair; read source (chopped assembly, HiFi or ONT reads); average coverage across aligned reads; "expensive" edge cost used in Floco; expected CN for most nodes, used in background read depth estimation; as well as method runtime and peak memory usage.

| Pair | Alignment source | Coverage | Expensive cost | Expected CN | Runtime (hh:mm:ss) | Memory (Gb) |
|---|---|---|---|---|---|---|
| CHM13-pang | Assembly | 20 | $-10^5$ | 1 | 01:03:30 | 22 |
| CHM13-pang | HiFi | 33 | $-10^5$ | 1 | 02:27:19 | 45 |
| CHM13-pang | ONT | 112 | $-10^5$ | 1 | 01:50:11 | 34 |
| HG01258-pang | Assembly | 7 | $-10^3$ | 2 | 00:40:30 | 35 |
| HG01258-pang | HiFi | 18 | $-10^5$ | 2 | 01:57:01 | 35 |
| HG01258-pang | ONT | 7 | $-10^5$ | 2 | 02:57:25 | 53 |
| HG01114-pang | Assembly | 7 | $-10^3$ | 2 | 01:29:00 | 138 |
| HG01114-pang | HiFi | 17 | $-10^5$ | 2 | 01:53:49 | 35 |
| HG01114-pang | ONT | 26 | $-10^5$ | 2 | 02:01:55 | 35 |
| HG01114-asm | Assembly | 17 | $-10^5$ | 1 | 00:10:36 | 2.4 |
| HG01114-asm | HiFi | 20 | $-10^5$ | 1 | 00:08:00 | 2.4 |
| HG01114-asm | ONT | 26 | $-10^5$ | 1 | 00:14:03 | 2.4 |
| Altus-asm | Assembly | 19 | $-10^5$ | 1 | 00:08:25 | 1.7 |
| Altus-asm | HiFi | 28 | $-10^5$ | 1 | 00:11:24 | 1.3 |
| Altus-asm | ONT | 65 | $-10^5$ | 1 | 00:22:08 | 3.1 |

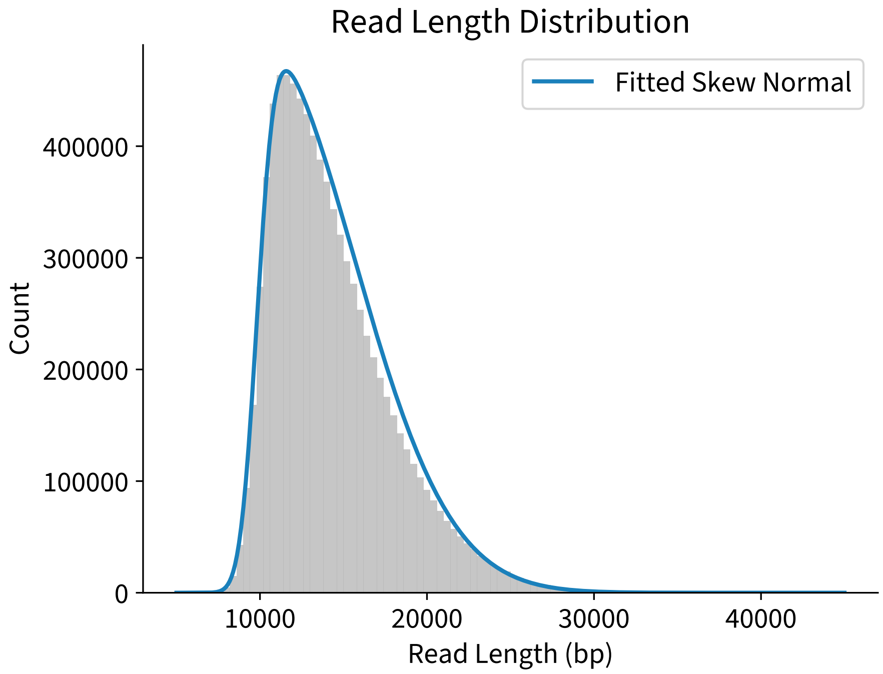

**Supplementary Figure 1. Read length distribution.** Distribution of HG01114 HiFi read lengths and the fitted Skew Normal distribution.

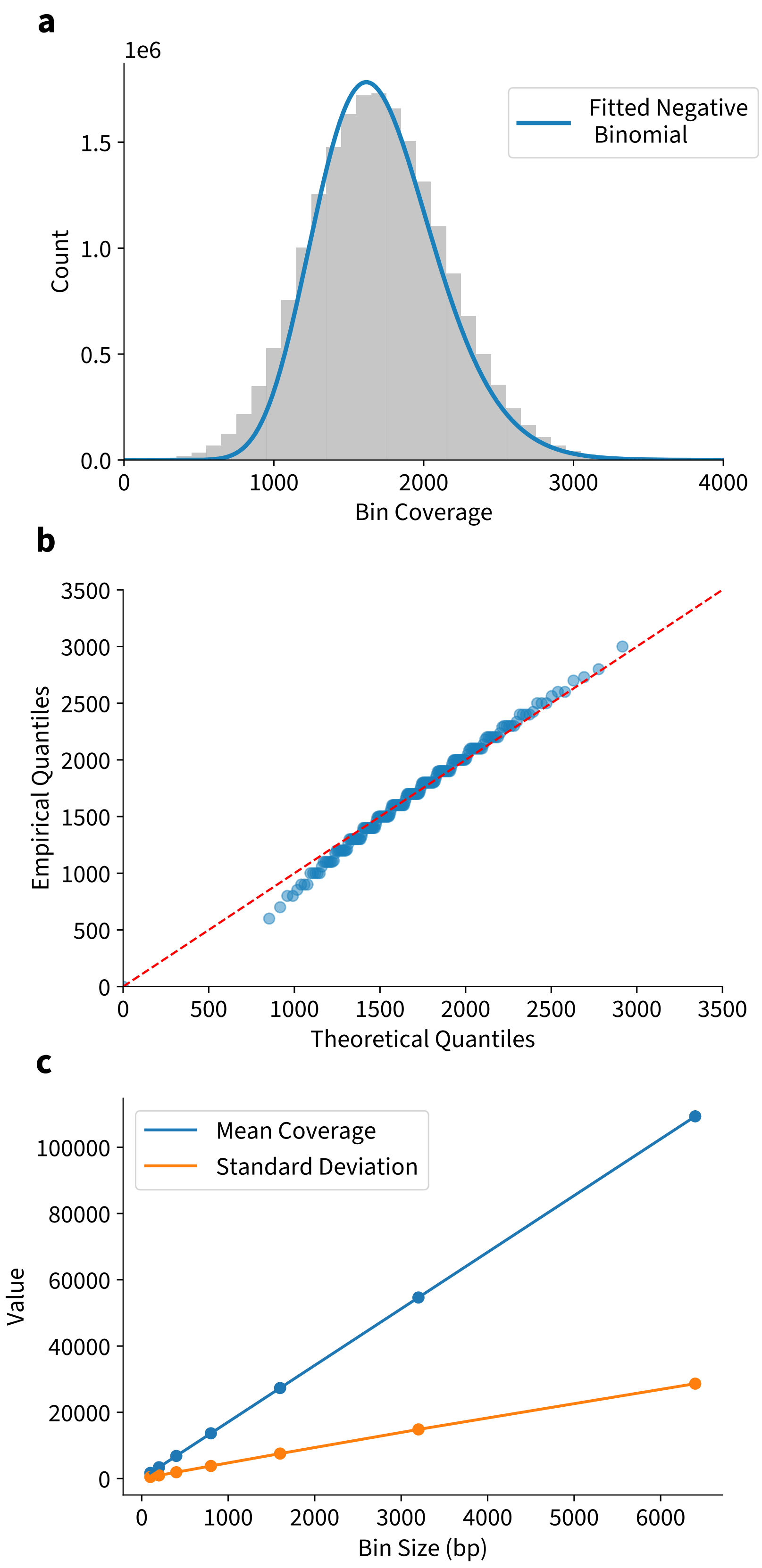

**Supplementary Figure 2. Bin coverage distribution. a**, Distribution of 100 bp bin coverages for HG01114-asm HiFi reads aligned against the HG01114 assembly graph, and the fitted Negative Binomial distribution. **b**, Corresponding theoretical and observed quantiles between 0.005 and 0.995 in 0.005 increment. Quantiles closer to 0 and 1 are not shown due to the presence of nodes with CN=0 and CN>1. **c**, Observed mean coverage (in blue) and standard deviation (in orange) for bin sizes from 100 bp to 6400 bp.

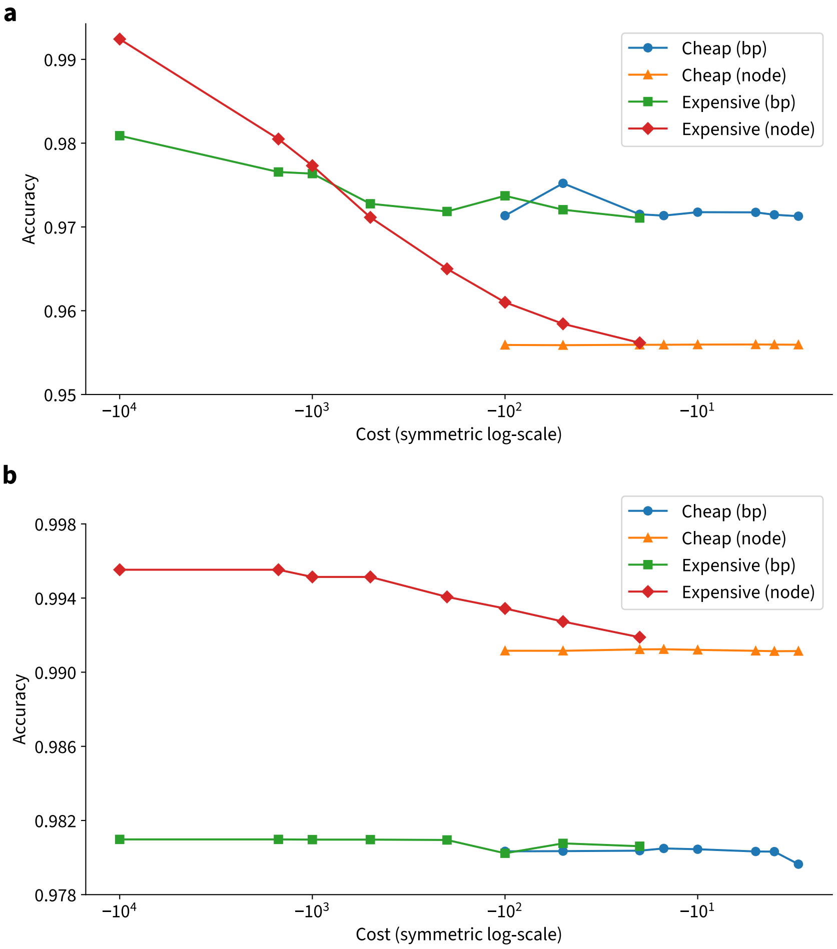

**Supplementary Figure 3. Floco accuracy depending on the superedge costs.** CN calling accuracy was evaluated for different cheap and expensive superedge costs using alignments from PacBio HiFi (**a**) or ONT (**b**) reads for the CHM13-pang pair. Accuracy was measured as the number of correctly assigned nodes as well as base pairs and displayed using a symmetric log-scale. To evaluate the accuracy based on the cheap costs, we used a fixed value of -10 for the expensive cost. Vice versa, we fixed cheap cost to -2 when evaluating expensive costs.

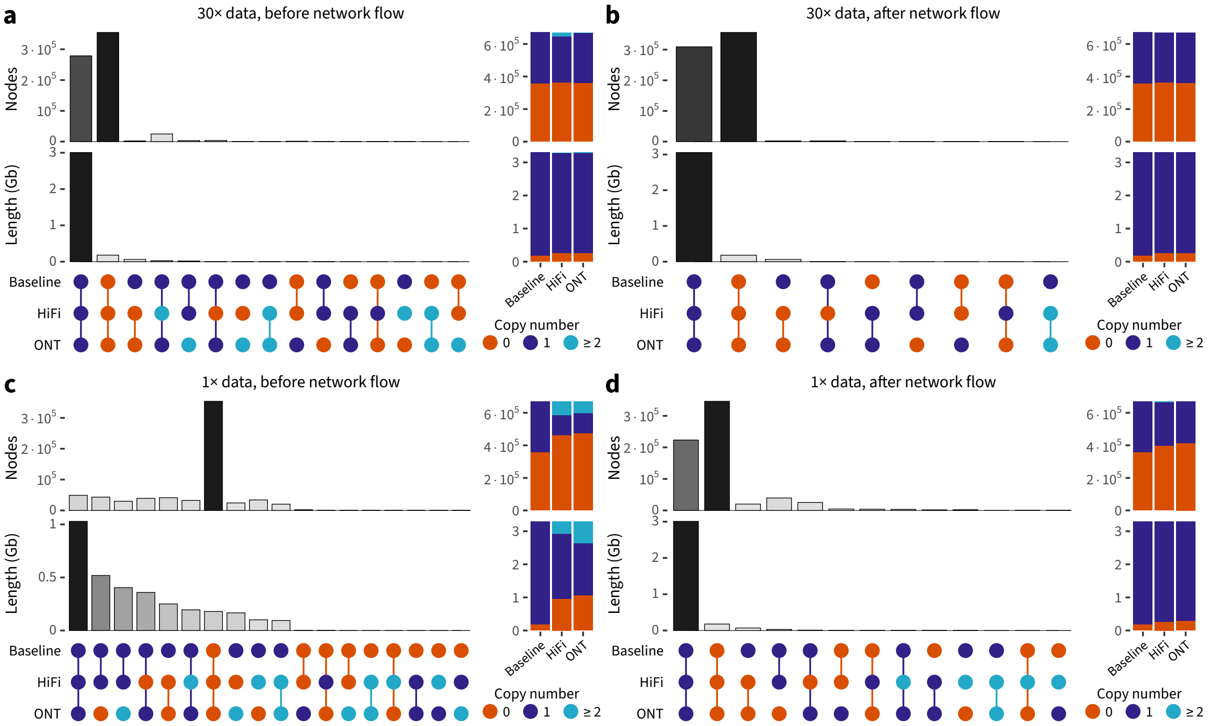

**Supplementary Figure 4. Copy number predictions at the CHM13 dataset with and without the flow network.** Extended UpSet plot with combinations of copy number values in the baseline (ground truth) CHM13 set, as well as Floco HiFi and ONT-based predictions. For each combination, corresponding node count and sum length are shown on top. The panels show CN predictions **a**, before network flow, 30× coverage; **b**, after network flow, 30× coverage; **c**, before network flow, 1× coverage; **d**, after network flow, 1× coverage.

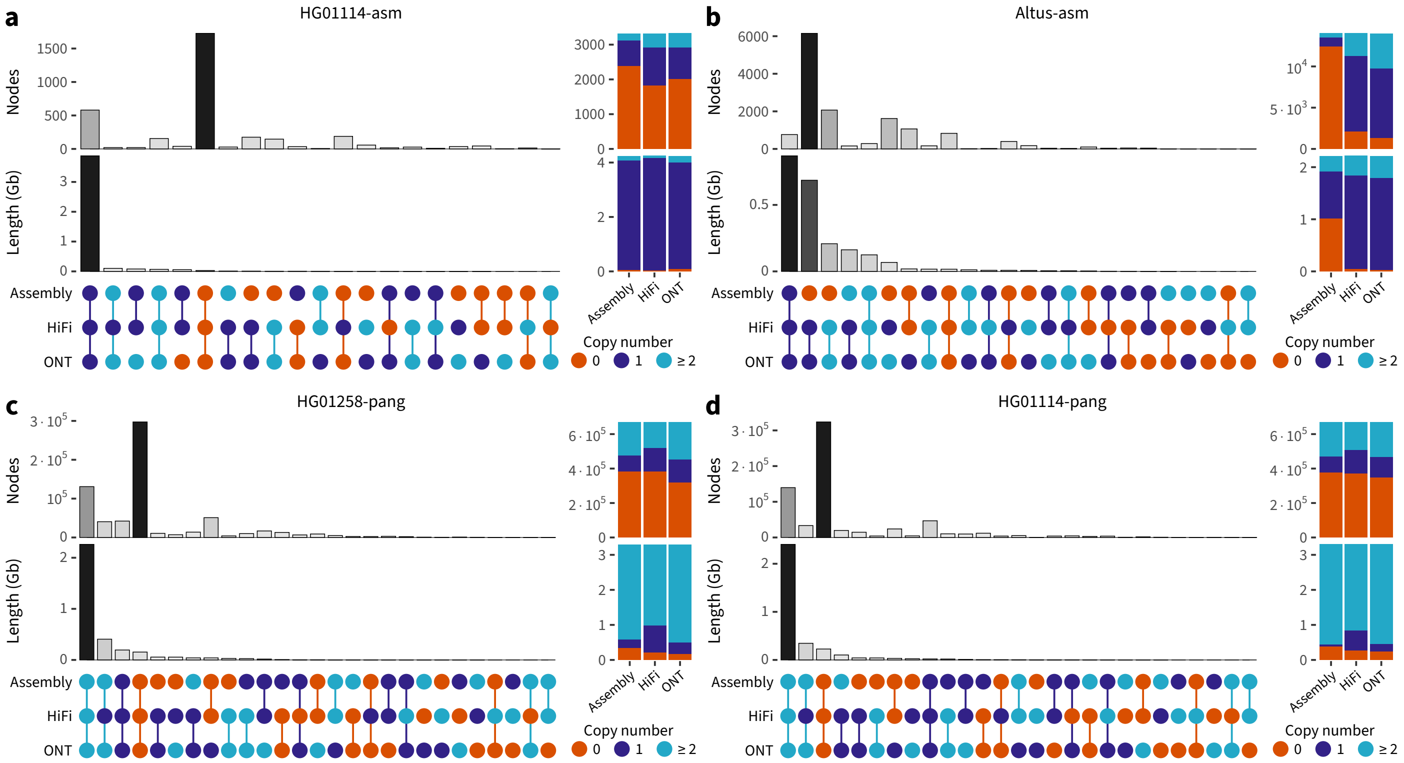

**Supplementary Figure 5. Concordance between copy number predictions for different sequencing technologies.** Panels show extended UpSet plots for different combination of copy number predictions based on the chopped assembly, PacBio HiFi and Oxford Nanopore (ONT) reads. Columns are sorted by sum length, all columns are shown. Total number of nodes as well as the corresponding sum length are shown on the right of every panel. Concordance is calculated for **a**, HG01114 sample mapped to its assembly graph; **b**, Potato Altus dataset mapped to its assembly graph; **c**, HG01258 sample mapped to the HPRC pangenome (in-sample); and **d**, HG01114 sample mapped to the HPRC pangenome (out-of-sample).

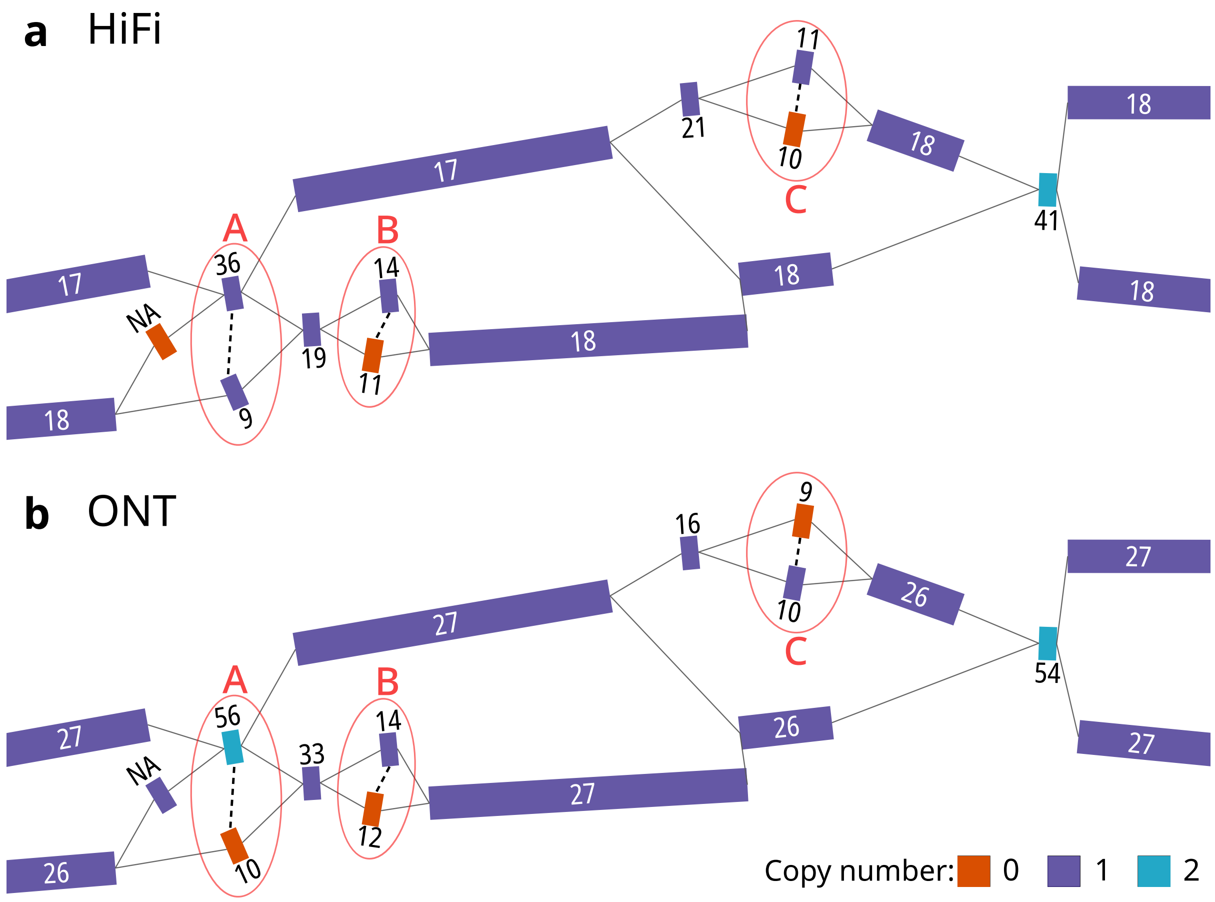

**Supplementary Figure 6. Copy number predictions at almost identical bubble arms.** strangepg visualization of part of the chromosome 6 with Floco CN estimations for the HiFi (**a**) and ONT (**b**) data for the HG01114 and the corresponding assembly graph. Nodes are colored by predicted CN (0: orange, 1: purple, 2: blue), numbers on the nodes show average read depth across the node. HiFi and ONT datasets have average coverage 20× and 26×, respectively. Three bubbles are highlighted with red circles and named A,B,C. All three bubbles contain almost identical pairs of nodes (denoted by dashed lines), with edit distances 1, 1 and 2 out of node sizes 22506, 11542 and 29542 for bubbles A, B and C, respectively. However, due to fluctuations in observed read depth, Floco predicted discordant copy number values for the two technologies at the bubbles A and C, still predicting concordant CN values at the bubble B. Node to the left of the bubble A contains no unique sequence and consequently has no associated read depth.

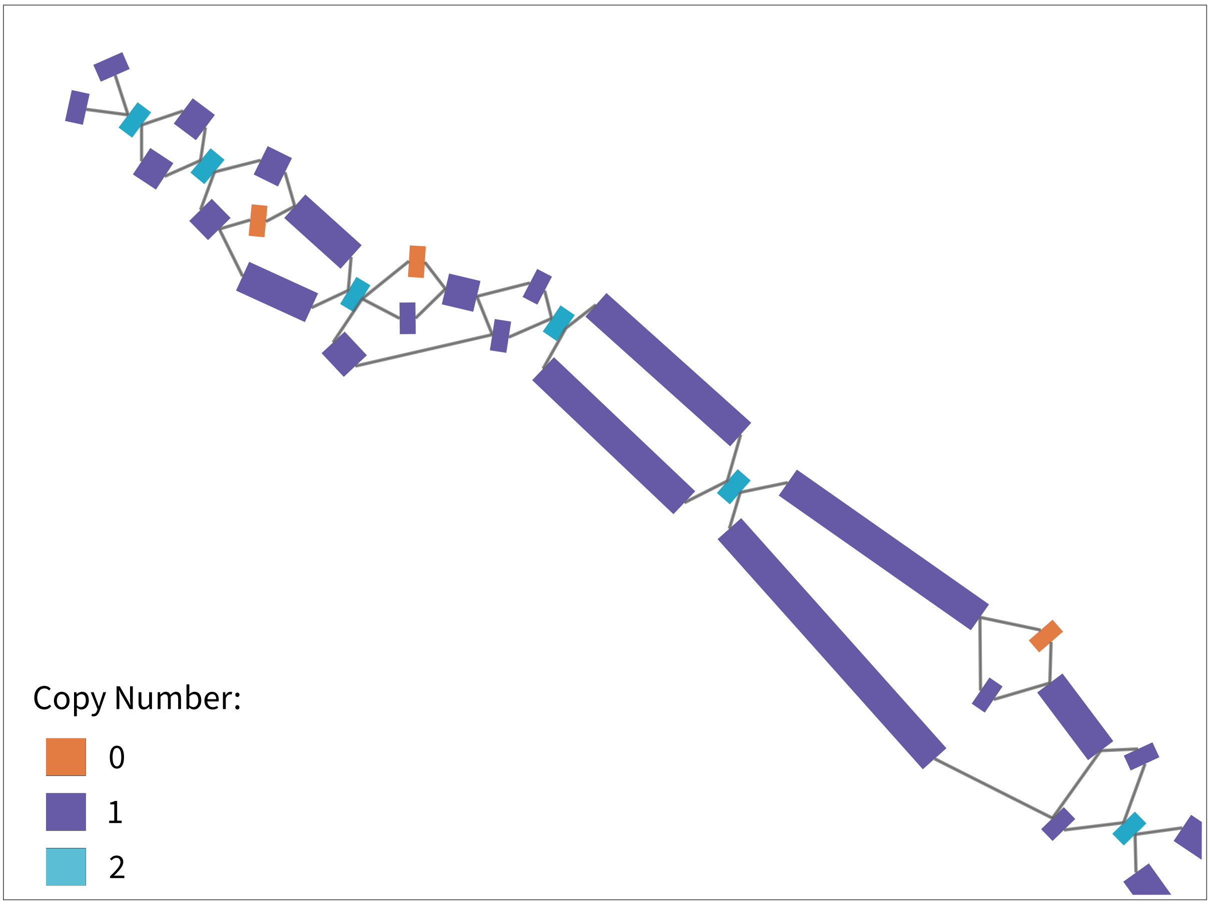

**Supplementary Figure 7. Example of flagged misassemblies.** strangepg visualization displaying 3 nodes with CN=0, which is, in theory, not expected when aligning HiFi reads from a sample against its own graph. In this case, these are misassemblies in the graph construction, which were manually confirmed, that Floco was able to successfully flag.

### 2 Supplementary Methods

#### 2.1 Graph preprocessing

During graph preprocessing, Floco clips nodes using a set of hierarchical rules in such a way that all transformed nodes do not overlap each other, and as little sequence is lost as possible. Specifically, we sort the graph edges in a descending order by overlap size $l_e$ and clip the overlapping sequence from one of the nodes unless both nodes have already been clipped on the previous iterations. For a given edge $e = (v, i, u, j)$ connecting node end $i \in \{0, 1\}$ of $v$ to node end $j$ of $u$, we go through a list of conditions and test each of them for both $(v, i)$ and $(u, j)$. Whenever a condition applies, we clip the node end in question and continue to the next edge. The conditions are written in terms of the node end $(v, i)$, but are equally tested for the node end $(u, j)$. In summary, we clip node end $(v, i)$ whenever:

- node end $(u, j)$ is already clipped (there may exist a third node that contains the same overlapping sequence);
- node end $(v, i)$ has more edges $\left|E^-_{vi}\right| + \left|E^+_{vi}\right| < \left|E^-_{uj}\right| + \left|E^+_{uj}\right|$ (in later iterations, a node with a bigger number of connected edges has a higher chance to be examined again, reducing the number of potential future clippings);
- node $v$ is longer: $l_v > l_u$ ($v$ is less likely to become fully clipped).

If none of the conditions applied, we clip the node that appeared later in the graph. After the full clipping procedure some nodes may be clipped from both sides such that the clippings overlap, and will effectively have zero or negative length. Such nodes will not possess read depth and their copy number will be estimated solely based on the copy number of the neighboring informative nodes.

#### 2.2 Benchmarking Floco

To generate "chopped" assembly datasets, we split linear assemblies into 20 kb reads without adding any new mutations or errors. Groups of reads were generated with the following steps: 2 kb, 5 kb, ..., 14 kb and 17 kb for CHM13-pang, HG01114-asm and Altus-asm cases, resulting in 20× coverage artificial datasets; as well as 5 kb, 10 kb and 15 kb steps for the HG01114-pang and HG01258-pang reads, producing 7× coverage datasets.

We then used GraphAligner `v1.0.19` [1] to map HiFi, ONT and assembly reads to their corresponding graphs. The following GraphAligner parameters were used:

- Potato Altus-asm (all technologies): `--bandwidth 15 --seeds-mem-count 100000 --seeds-mxm-windowsize 10000`

- Assembly and HiFi reads, mapped to the HPRC pangenome (CHM13-pang, HG01258-pang, HG01114-pang): `--bandwidth 15 --multimap-score-fraction 0.99 --precise-clipping 0.9 --min-alignment-score 2000 --seeds-mem-count 100000 --seeds-mxm-windowsize 10000`
- ONT reads, mapped to the pangenome (CHM13-pang, HG01258-pang, HG01114-pang): `--bandwidth 15 --multimap-score-fraction 0.99 --precise-clipping 0.75 --min-alignment-score 2000 --seeds-mem-count 100000 --seeds-mxm-windowsize 10000`
- Assembly reads for HG01114-asm: `--bandwidth 15 --seeds-mem-count 100000 --seeds-mxm-windowsize 10000 --multimap-score-fraction 0.99 --min-alignment-score 2000`
- HiFi and ONT data for HG01114-asm: `--seeds-mxm-windowsize 5000 --seeds-mxm-length 30 --hpc-collapse-reads --seeds-mem-count 10000 --bandwidth 15 --multimap-score-fraction 0.99 --precise-clipping 0.85 --min-alignment-score 5000 --discard-cigar --clip-ambiguous-ends 100 --overlap-incompatible-cutoff 0.15 --max-trace-count 5`

To calculate Floco accuracy for the CHM13 data, we assigned ground truth CN=1 for nodes that have chromosome names in their SN tags, otherwise, nodes were assigned CN=0. Correspondingly, nodes with ground truth CN=1 and Floco-predicted CN=1 were considered true positive (TP) events. Ground truth CN=1 and any other CN predicted by Floco were considered to be false negative (FN), even when Floco could predict positive CN other than CN=1. Correspondingly, true negative (TN) and false positive (FP) events have ground truth CN=0 and Floco-predicted CN=0 and CN≠0, respectively. We then calculated precision and recall based on the number of TP, FP and FN events, as well as their sum node length. Simple accuracy was calculated as the fraction (and fraction weighted by length) of nodes with matching ground truth–Floco CN values, which is equivalent to (TP + TN) / (TP + TN + FP + FN).

To calculate Floco concordance across different data sources, we calculated the frequencies of all combinations of CNs. For the plots, we merged all CN predictions ≥2 to reduce the total number of events. For concordance value calculation, described in the main text, we merged all CN predictions ≥4 as it becomes increasingly difficult to distinguish between high CN values (e.g. 5 and 6), while their phenotypical impact becomes less distinctive. Node-based concordance for the assembly graphs was performed using informative nodes only since fully clipped nodes are fully overlapped by neighbouring nodes and no read depth and possess no independent read depth information.

We used Locityper `v1.2` [2] align module to find global alignments between various nodes in the graph (Supplementary Figure 6). All graph visualizations were performed using strangepg [3].

[3] Bonnet, K. & Marschall, T. strangepg. https://github.com/qwx9/strangepg.
